# Properties of compostite feedback-feedforward pulse generating motifs

**DOI:** 10.1101/074377

**Authors:** Bharat Ravi Iyengar

## Abstract

Negative feedbacks and incoherent feedforward loops are known to give rise to a pulse in response to a step change in the input. In this article, I present a study of composite motifs made of coupled feedback and feedforward loops, acting via different regulatory mechanisms. In these motifs, the effect of input and output on the controller is realized via either an AND-gate or an OR-gate. Using a simplistic model of gene expression and a common parameter set, I have studied the effect of global parameter variation on the dynamic and steady-state properties of different motifs, in response to a step change in the input. These metrics include steady state gain, response time, overshoot, peak time and peak duration. For the motifs with a negative feedback component, it can be seen that AND-gated motifs show a “feedforward-like” property whereas the OR-gated motifs show a “feedback-like” property. Motifs with a positive feedback component show hypersensitivity of gain, to parameters. Overshoot correlates negatively with peak time whereas peak duration correlates concavely with peak time, a property that is also observed for uncoupled feedback and feedforward motifs. This indicates that this relationship between overshoot, peak duration and peak time, seems to be a universal property of pulse-generating motifs.

## Introduction

Gene expression is a highly regulated process in which the regulators affect the concentration and activity of the gene product by controlling different steps of its synthesis^[1]^. The interactions between different genes in the genome constitutes the gene regulatory network (GRN), in which genes represent nodes and the regulatory interactions represent the edges. Since the regulatory interactions have a direction and can have a positive or a negative effect on the target node, the GRN is a signed and directed network. Some patterns of subnetworks are much more commonly found in a realistic network (such as GRN) compared to what would be expected in a randomized network. These patterns are known as network motifs and different network motifs usually have unique functional properties^[2,3]^. Feedback and feedforward loops are among the best studied network motifs in the GRN ^[3,4]^ and some of their subtypes are also well known control mechanisms studied in control theory^[5]^.

In a feedback loop, a node regulates itself either directly or indirectly. There are two kinds of feedback loops – positive and negative. The product of all the signs of individual edges gives the overall sign of the loop. In other words, if the overall effect of the node (gene) on itself is positive then the motif is known as a positive feedback loop (and vice versa).

In a feedforward loop, a node regulates another node via two parallel paths. If the overall signs of these paths are the same then the motif is known as a coherent feedforward loop or else, an incoherent feedforward loop^[6]^.

Both the negative feedback and the incoherent feedforward loops can give rise to dynamic pulses in the expression of the output gene, in response to a step change in the input. In some cases, these motifs can also lead to a perfect adaptation^[7,8,9]^ i.e. the output steady state level after the step change is same as the initial level, while the transient levels are different. However, a simple feedback loop which does not have a perfect integral feedback does not lead to perfect adaptation (^[10]^). Under certain conditions, the incoherent feedforward motif can also exhibit fold change detection, a phenomenon in which both the dynamics and the steady state of the output, are only dependent on the fold difference between the input levels^[8, 11]^.

Though the properties of feedback and feedforward motifs have been studied in great detail, the properties of a generalized system that has coupled feedback and feedforward loops, to my knowledge, have not been analysed. An exhaustive list of examples of such motifs is not documented in the scientific literature but there are a few examples that suggest that these motifs may be prevalent in the GRN and cell-signalling networks. Tolllike receptors are known to activate the production of both the inflammatory and anti-inflammatory cytokines that in turn, mutually repress each other^[12]^. Nodal and lefty pathways are known to form a negative feedback loop that is critical for mesoderm development^[13, 14]^. In zebrafish, the nodal agonist, *squint* and the antagonist, *lefty* are targeted by a common upstream regulator – miR-430^[15]^. Another miRNA, miR-449 regulates both Notch and Delta^[16]^, which mutually repress each other within a cell (*cis*) and activate each other when present in different cells (*trans*).

In this study, I have focussed on regulatory motifs in which the input signal has a positive effect on expression of the output and the controller genes whereas the controller has a negative effect on the expression of the output. Furthermore, the output gene product can either repress or activate the controller. If the action of the output on the controller is removed then this motif reduces to a type-1 incoherent feedforward loop. Similarly, if the action of the input on the controller is removed then this motif reduces to a feedback loop (positive or negative depending on the action of the output on the controller). Likewise, removing the effect of the output on the controller will result in another incoherent feedforward loop (Fig. 1).

**Figure 1:**
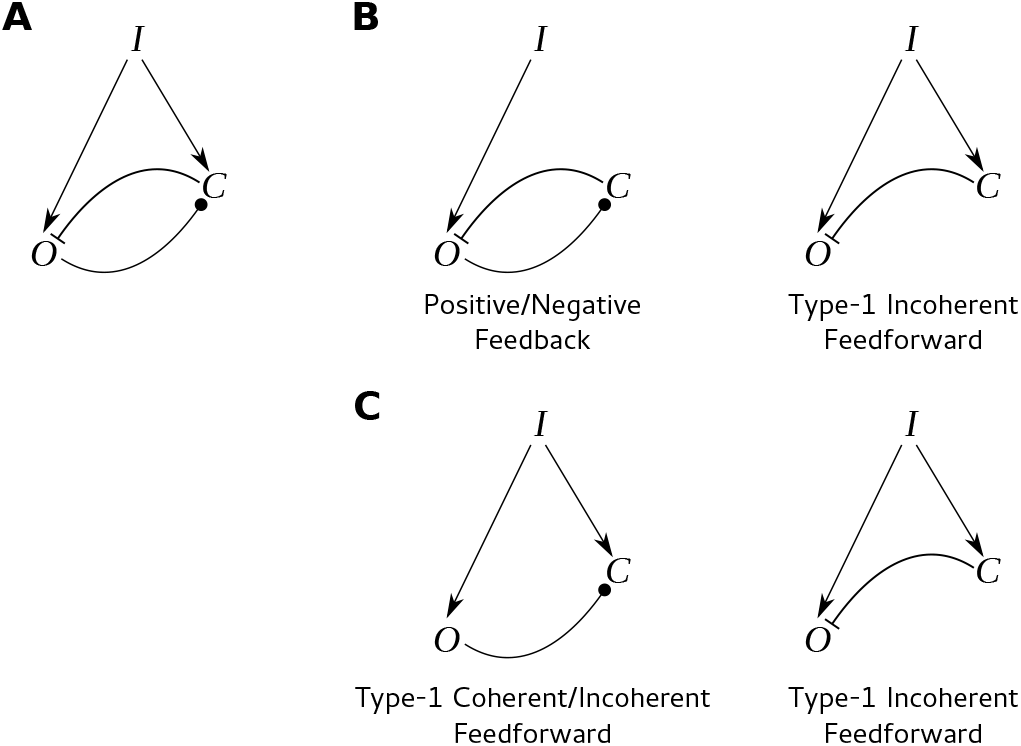
The composite feedback-feedforward motif (A) which can be decoupled to give rise to a feedback and a feedforward motif (B) or two feedforward motifs (C). Here, I denotes the input, *O* denotes the output and *C* denotes the controller. Arrowheads denote activation, flat ends denote repression and round end denotes any type of regulation.

I have analysed different steady-state and dynamic properties of composite feedback-feedforward motifs using a mechanism based mathematical model and well defined metrics. These metrics include steady-state gain, response time, overshoot, peak time and peak duration. The combined effect of positive regulation by the input and the output, on the controller can be either additive (OR-gated) or multiplicative (AND-gated). Both these cases, along with different modes of regulation by the controller, have been modelled and studied simultaneously. Additionally, I have also modelled a case in which the output has a negative effect on the controller i.e. the output and the controller mutually repress each other. All the models use a common set of parameters, obtained from values reported in published studies. Random multivariate sampling was used to study the global sensitivity of different motif properties to parameter changes. This study reveals that the AND-gated motifs exhibit “feedforward-like” properties (based on the metrics) whereas the OR-gated motifs exhibit “feedback-like” properties. This study also reveals that in these composite motifs the pulse dynamics follows a previously observed pattern with feedback and feedforward loops – the overshoot correlates negatively with peak time and the peak duration correlates concavely with peak time^[10]^. This suggests that this property may be a universal feature of these pulse-generating motifs.

## Methodology

### Description of the models and the metrics

All the modelled motifs have four components (variables) – output mRNA (*r*_*o*_), controller mRNA (*r*_*c*_), output protein (*p*_*o*_) and controller protein (*p*_*c*_). The input is modelled as a parameter instead of a variable. The input always activates the formation of output and controller mRNAs. Also, the output protein always regulates the controller at the level of transcription. Four different modes of regulation are considered for the effect of controller on the output but the net effect of the controller on the output is always negative. If the output protein activates the controller then the resultant feedback within the composite motif would be negative and if it represses the controller, then the feedback would be positive (also referred to as “double-negative”). The controller can either repress the formation of the output mRNA or protein, or promote their degradation. The combined effect of input and output on controller can be multiplicative (AND) such that both the input and the output are necessary for regulation, or additive (OR) in which either of them can regulate the output. Since the combinatorial effect of a negative (output) and a positive (input) regulator on transcription of a gene (controller) is usually multiplicative, there are no OR-gated motifs when the feedback component is positive. All these combinations yield twelve different motifs (Fig. 2). These motifs are abbreviated using a code which would be used to refer to these motifs, hereafter. The first two letters of the code denote the mode of regulation – rf: RNA formation, rd: RNA degradation, pf: protein formation and pd: protein degradation. The last letter refers to the nature of the feedback component – n for negative and p for positive. The uppercase letters in-between denote the nature of the combinatorial effect – AND/OR.

**Figure 2:**
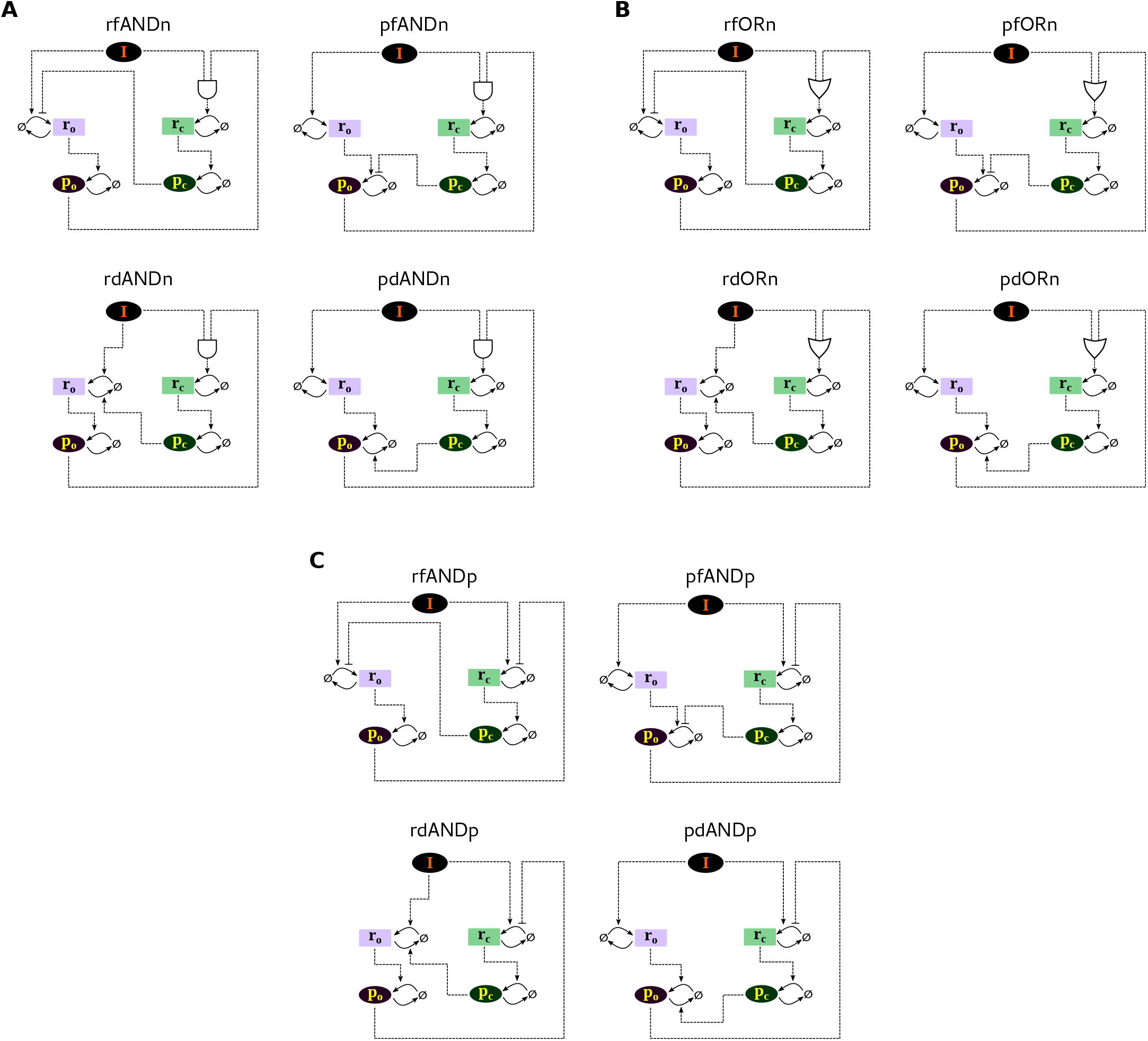
Different composite feedback-feedforward motifs analysed in this study. Panels A and B show AND-gated and OR-gated motifs with a negative feedback component. Panel C shows the composite motif with a positive feedback component. Each panel has four diagrams depicting the four modes of regulation (the abbreviations in the headings are explained in Methodology). In all the diagrams, input, output mRNA, controller mRNA, output protein and controller protein are denoted by *I* (black oval), *r*_*o*_ (lilac box), *r*_*c*_ (light-green box), *p*_*o*_ (purple oval) and *p*_*c*_ (dark-green oval), respectively. ø denotes the cellular pool of nulceotides and amino acids which is assumed to be infinite. In the regulatory edges (dashed arrows), arrowheads denote activation whereas flat ends denote repression. Solid arrows denote conversion.

The responses of these motifs were studied, for a step change in the input. The initial conditions were set equal to the steady state value of the system corresponding to a given value of input, *I*_1_. The input was changed to a higher value (step up), *I*_2_ and the dynamics of the response of the output protein was studied using different metrics (Fig. 3, Equations 1–5). These metrics are defined in our previous work^[10]^; I shall provide a brief description of these motifs, here.

**Figure 3:**
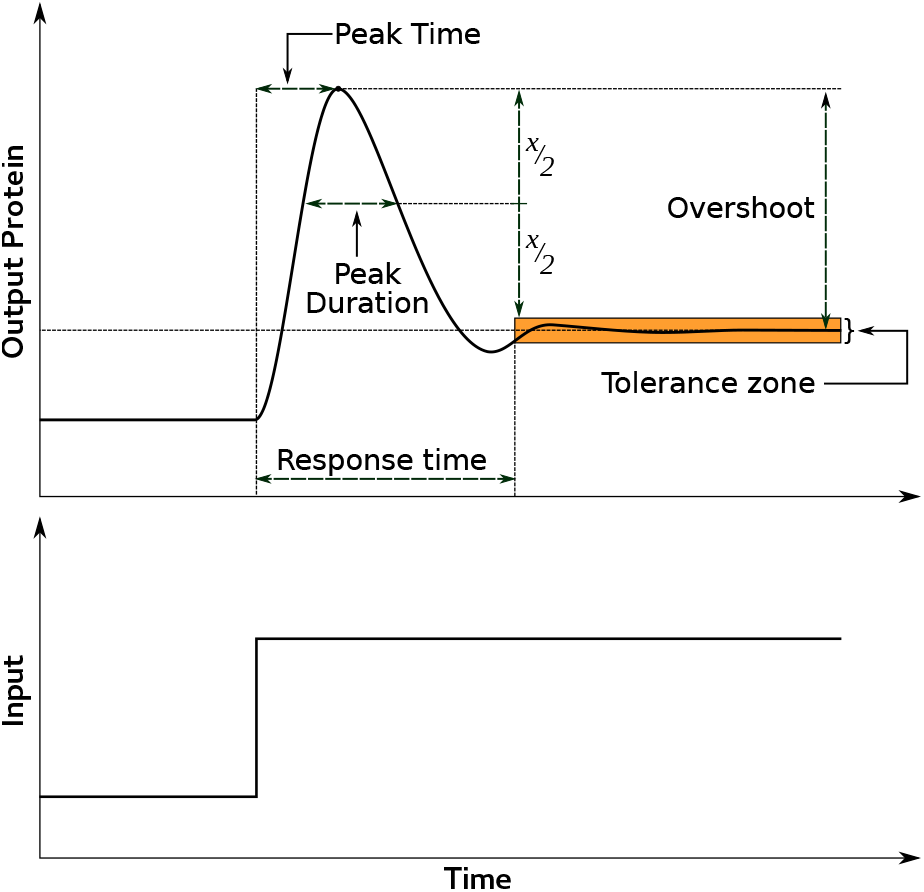
Illustration of the metrics defined in Equations 1–5 on a dynamic response curve of output protein (*p*_*o*_). Steady state gain is not shown in this illustration.

Steady-state gain is the ratio of the change in steady state concentration and the initial concentration of output protein.

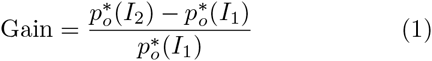

Here 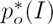 refers to steady state value of *p*_*o*_ for a given input *I*.

Response time (*t*_*r*_) is the time required for *p*_*o*_ to converge to a tolerance region defined as ±1% around 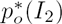.

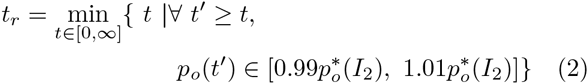

Overshoot is the difference between maximum dynamic value of *p*_*o*_ and the final steady state 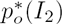. Overshoot is normalized to 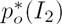 to obtain a per-unit overshoot and all further references to overshoot mean the per-unit overshoot.

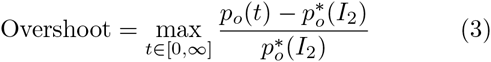

Peak time (*t*_*p*_) is the time required to reach the maximum value, normalized to response time.

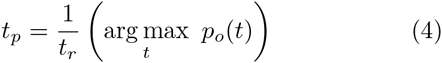

Peak duration is the width of the peak at half of its height above the upper bound of the tolerance zone, normalized to the response time.

**Table 1:**
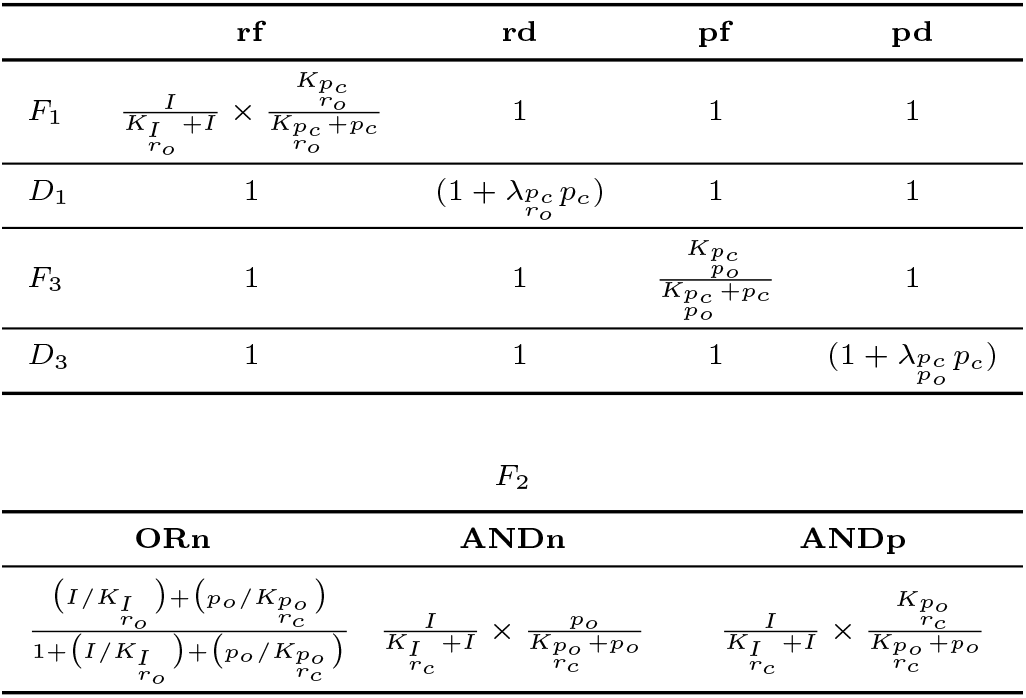
Details of the F and D functions used in the ODEs describing the mathematical model of the composite motifs (equations 6–9).

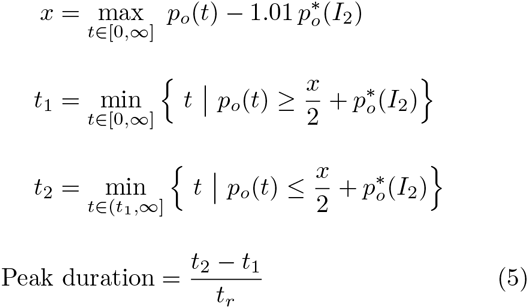

To study the global sensitivity of these metrics (equations 1–5) towards different parameters (except input), the metrics were calculated for 10000 different parameter sets that were randomly sampled around a basal value; this resulted in a distribution of these metrics.

### Mathematical model of the motifs

The mathematical models for the different motifs are described by a system of coupled non-linear ordinary differential equations with the concentrations of *r*_*o*_, *r*_*c*_, *p*_*o*_ and *p*_*c*_ as dependent variables (for convenience the concentrations have same symbol as the name of the species, as in the equations 1–4. Usual convention is to use square brackets). As previously mentioned, the input (*I*) was considered a parameter. The equations for different motifs have the following general form:

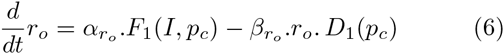

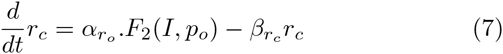

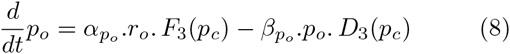

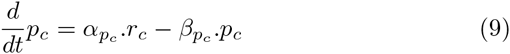

The functions, *F*_1_, *F*_3_, *D*_1_ and *D*_3_ represent the different modes of regulation by the controller protein whereas the function, *F*_2_ represents the combinatorial regulatory effect of input and output on the controller. The effect of a regulator on its target has been modelled using a saturating (Hill-like) function. The details of the F and *D* functions are shown in Table 1.

### Parameter sampling and simulation

A basal set of parameters was obtained from data available from published studies (Tab. S1). 10000 parameter sets were obtained by random multivariate sampling in an interval with lower and upper limits as 1/5 and 5 times the basal value, respectively. The sampling was done in a log-transformed range for a uniform representation of both higher and lower values. The initial (*I*_1_) and final (*I*_2_) value of input, were arbitrarily chosen as 4000 and 8000 molecules per cell and the volume of the cell is assumed to be 2.25 × 10^-12^*l*^[17]^. The steady state values were calculated using analytical expressions. For the step up simulations, the initial values were set as the analytical solution of steady state corresponding to *I*_1_. Then, the input was changed to *I*_2_ and the ODEs (equations 6–9) were numerically integrated using Matlab ode15s solver to obtain the temporal dynamics.

## Results and Discussion

### ANDn, but not ORn, motifs show perfect adaptation

Gain (Eqn. 1) for different motif systems, was calculated using analytical expressions for steady state, with the 10000 different parameter sets. It can be observed that AND-gated motifs with negative-feedback component (ANDn) showed perfect (or near perfect) adaptation which is reflected by a sharp peak of the gain distribution at 0 (Fig. 4, first column). Although the OR-gated motifs with negative-feedback component (ORn) did show this behaviour for a few parameter combinations (~ 2% cases of ORn compared to ~ 40% ANDn cases that showed gain ≤ 0.05), their mean gain (and the peak) was around 0.4, indicating that these motifs do not usually exhibit perfect adaptation. For some parameter sets both ANDn and ORn motifs also showed a negative gain (~ 13% of ANDn and ~ 0.3% of ORn cases) which indicates a net decrease in the output with an increase in the input. Overall, there was no significant difference between the distributions for different modes of regulation. To understand the net effect of the input on the gain, the steady state values of the output protein were plotted for different values of input, for 200 randomly chosen parameter sets (Fig. 5). The slope of curves (input-output characteristics) between two points is an indicator of gain. It can be observed that input-output characteristics of ANDn motifs show a saturating behaviour and in some cases, non-monotonicity. Saturation indicates that these cases show adaptation for different values of input. Nonmonotonic curves suggest that, for these parameter sets, the motif will show a positive gain in a certain range of input and a negative gain in another. A few cases of ORn motifs also have saturating input-output characteristics which again suggests that for a small number of cases ORn motifs can show perfect adaptation. However for most cases, the input-output characteristics for ORn motifs are monotonically increasing. It is apparent that most of the cases which showed saturation and non-monotonicity, were hypersensitive at low values of input. Considering our previous observations with uncoupled feedback and feedforward motifs^[10]^, it appears that the steady state properties of ANDn motifs have a more “feedforward-like” nature whereas those of ORn motifs have more of a “feedback-like” nature.

**Figure 4:**
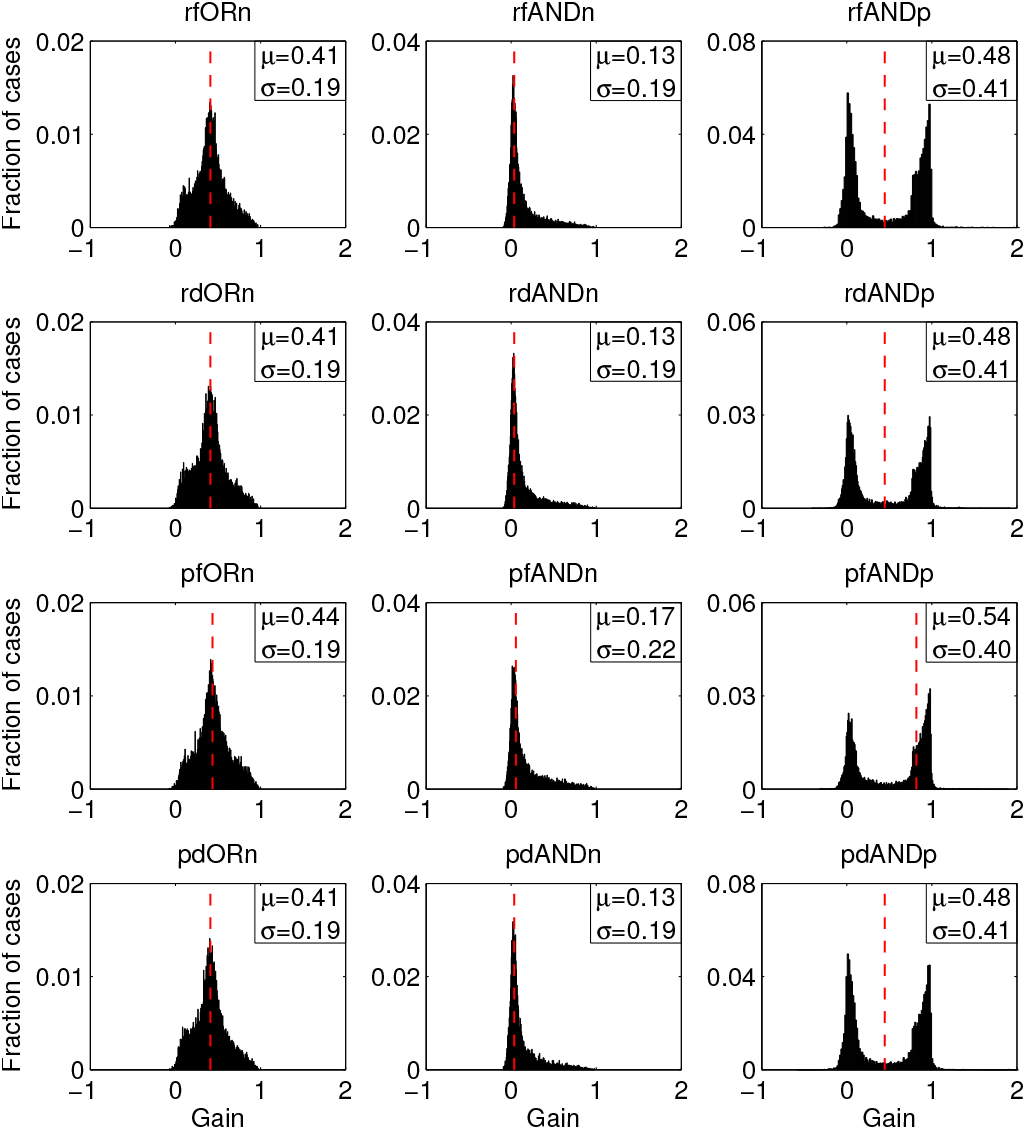
Steady state gain distributions for the different composite motifs. *µ* and *σ* indicate the mean and standard deviation of the corresponding distributions. The dashed vertical red line denotes the value of gain for the basal parameter set.

**Figure 5:**
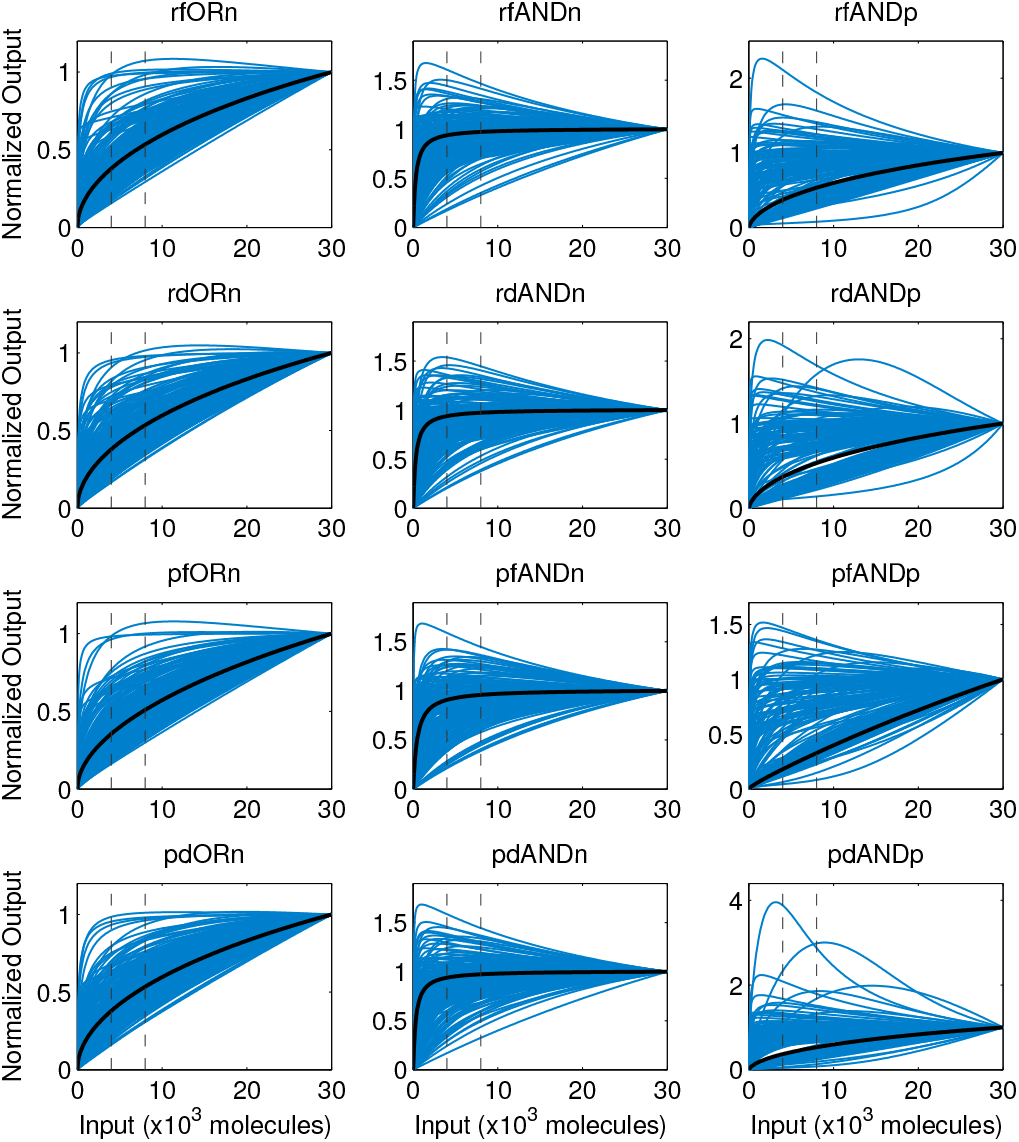
Input-output characteristics of the different composite motifs. The output values at each value of input, are normalized to the value at the maximum value of input (30 × 10^3^). Black line represents the characteristics for the basal parameter set. Dashed vertical lines (left and right) denote the values of *I*_1_ and *I*_2_ used in the analyses, respectively.

### Gain is hypersensitive to parameters, for ANDp motifs

Composite motifs with positive-feedback component (ANDp) showed a bimodal distribution of steady state gain with two sharp peaks at 0 and 1 (Fig. 4, last column). This indicates that the these motifs exhibit hypersensitivity of gain towards different parameters such that the output protein either perfectly adapts (gain=0) or shows no regulation at all (gain=1). ANDp motifs showed negative gains for a few cases (~ 9%); for a small number of cases it even showed gains>1 (~ 1.3%). As in case of ANDn and ORn motifs, there were only minor differences between the gain distributions for different modes of regulation. The input-output characteristics for ANDp motifs also showed non-monotonicity and saturation, in some cases. However, there were also cases in which the output was not hypersensitive to input unlike the case of ANDn motifs. Some curves were also concave, which indicates a gain>1 (Fig. 5, last column). Since the output and the controller are mutually repressive, I wanted to see if their gain distributions are reciprocal. A scatter plot between steady state gain for output-protein and that of the controller-protein (Fig. 6) revealed that these quantities are indeed reciprocal in the extreme regions (around the 0 and 1 peaks). Overall, there is an apparent negative linear correlation between these quantities. Interestingly, even ORn motifs show a negative linear correlation between steady state gains of output and controller but ANDn motifs do not.

**Figure 6:**
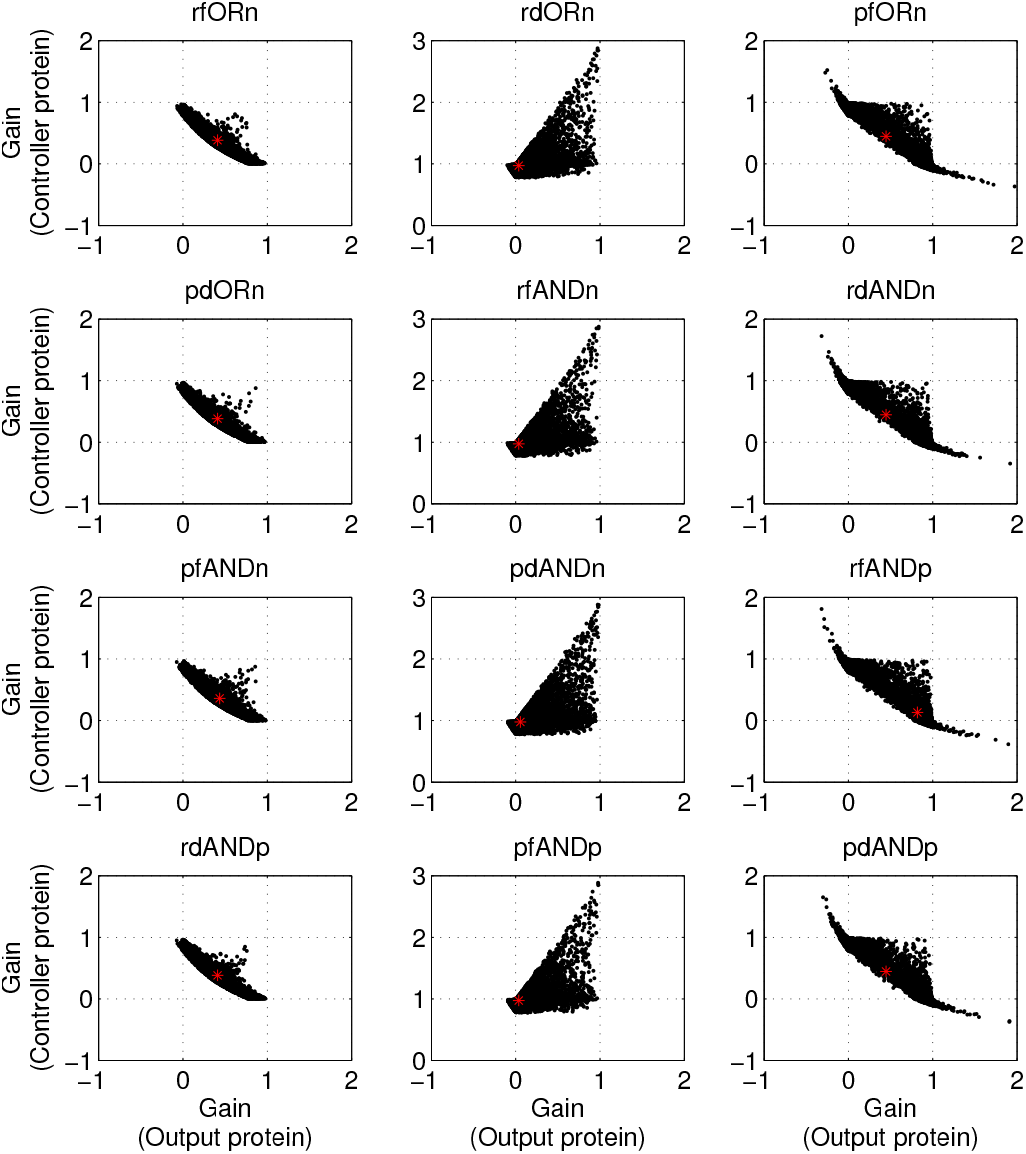
Scatter plot between steady state gain of output and controller proteins, for different composite motifs. The red asterisk denotes the value corresponding to the basal parameter set.

The gain distributions for all the twelve motifs retained their overall shape for a broader range of parameter variation – 1/20 to 20 times the basal value (Fig. S1).

### Regulation by protein degradation has the fastest response compared to other modes of regulation

Response time and other dynamic properties were calculated after performing the step-up simulation. From the distributions shown in Fig. 7, it could be observed that motifs in which the controller acts via promoting the protein degradation (pd motifs), have the fastest responses i.e. low response time, compared to other modes of regulation. The response time distributions for the pd motifs were also narrower (low standard deviation) compared to that of other motifs. Even in the case of uncoupled feedforward and feedback motifs, regulation by protein degradation is faster than other modes of regulation (Fig. S2). However, the mean and the variance of response time distributions of the feedback motifs were lower than that of the feedforward motifs, for the corresponding modes of regulation.

**Figure 7:**
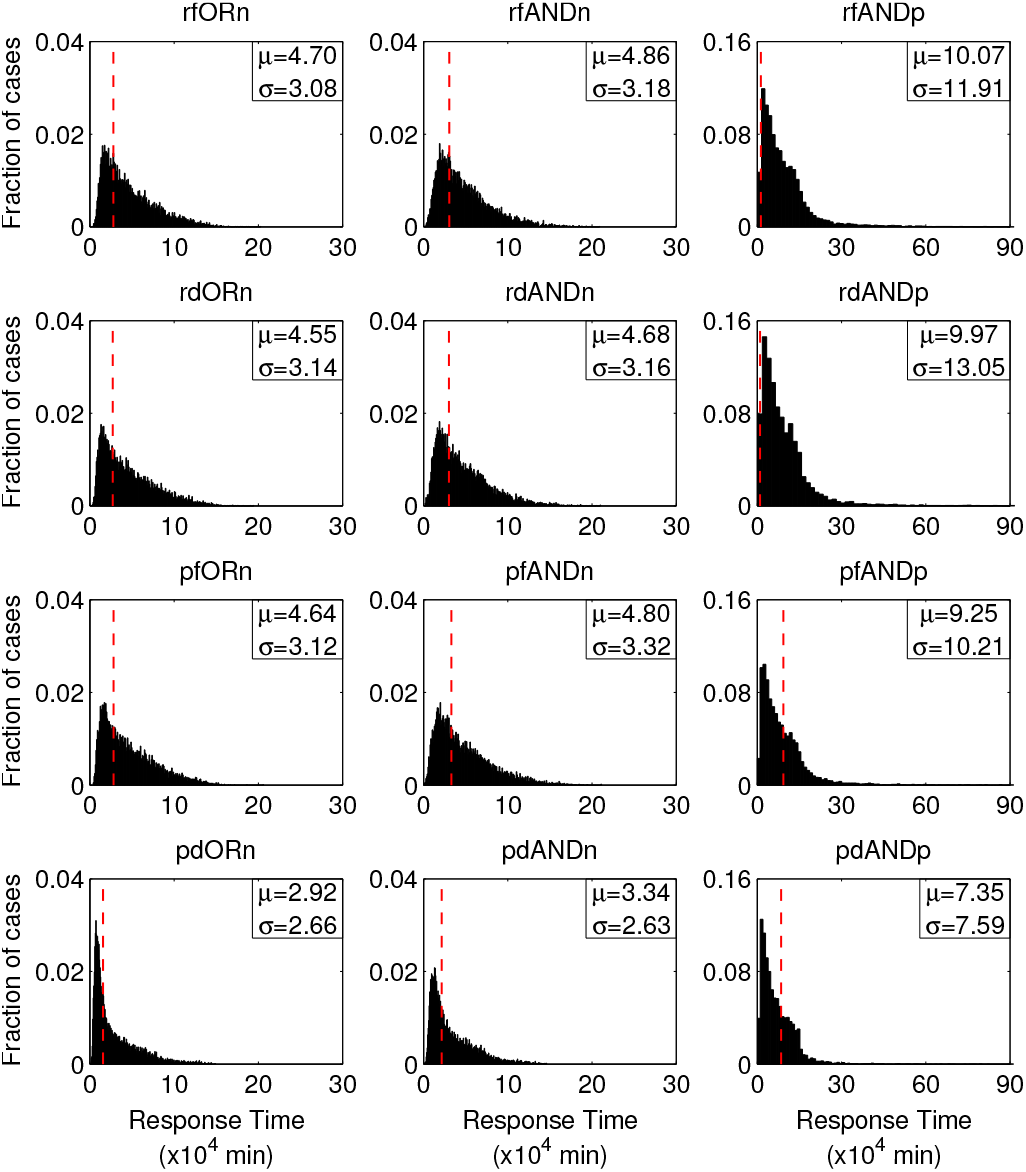
Response time distributions for different composite motifs. *μ* and *σ* indicate the mean and standard deviation of the corresponding distributions. The dashed vertical red line denotes the value of gain for the basal parameter set. The few values exceeding 90×10^4^ (<0.5%) in case of ANDp motifs, are not shown.

### ANDp motifs have higher response times compared to ANDn and ORn motifs

ANDp motifs have higher mean response times as well as a broader distribution relative to ANDn and ORn motifs of corresponding modes of regulation (Fig. 7). In fact, it can be noticed that even the co-efficient of variation (standard deviation/mean) is much higher than that of the other motifs. Moreover, some few isolated cases showed extremely high response times (more than 6 standard deviations from the mean). However, even for ANDp motifs, regulation via protein degradation has a lower mean and standard deviation (also the co-efficient of variation) of response time.

### ANDn and ANDp motifs have higher mean overshoot and a broader distribution compared to ORn motifs

A system is said to exhibit a peak when it has non-zero overshoot. In more than 67% cases (out of 10000 parameter sets), the ORn motifs exhibited peaks (Fig. 8) with the pdORn motif showing the highest peaking propensity (~ 92%). The ANDn motifs also showed very high propensity to generate peaks; ~ 90% of cases showed peaks for all modes of regulation except pfANDn for which it was ~ 80%. However, the ANDp motifs showed much less peaking propensity; only ~ 45% cases of pfANDp and ~ 50% cases of other motifs showed peaks. Since the generic structure of the ANDp motifs is symmetric with respect to output and controller (as they are mutually repressive), I expected that many of the cases of ANDp that did not show overshoot of output protein, might instead show an overshoot of the controller protein. This indeed turned out to be the case (Fig. S3). More than 93% of cases that did not show overshoot of output protein showed an overshoot of the controller protein while the rest did not show overshoot for both the proteins. Like steady state gain, the overshoot also seems to follow this mutual exclusivity between output and controller and hypersensitivity to parameters.

**Figure 8:**
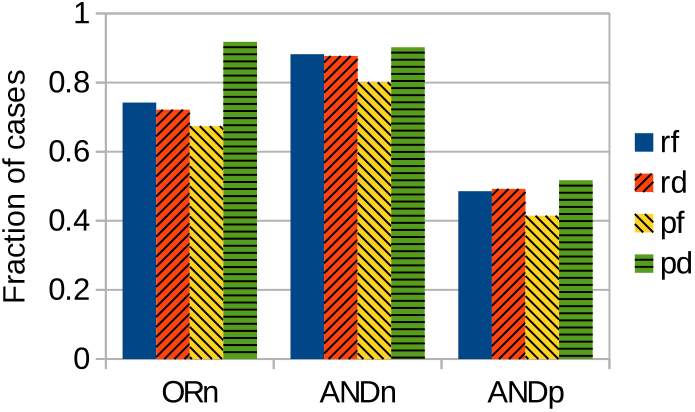
Peaking propensity for different composite motifs. Colour/pattern of bar fills denote different modes of regulation (rf: solid blue, rd: orange; 45^0^ diagonal hatch; pf: yellow, -45^0^ diagonal hatch and pd: green, horizontal hatch)

For the cases that exhibited peak (for output), overshoot, peak time and peak duration were measured. The distribution of overshoot (Fig. 9) for the ANDn and the ANDp motifs had a higher mean and standard deviation, than that of the ORn motifs with corresponding mode of regulation. However, the peak time distributions showed the opposite trend; the ANDn and the ANDp motifs had a lower mean and narrower distribution of peak time, compared to the ORn motifs (Fig. 10).

**Figure 9:**
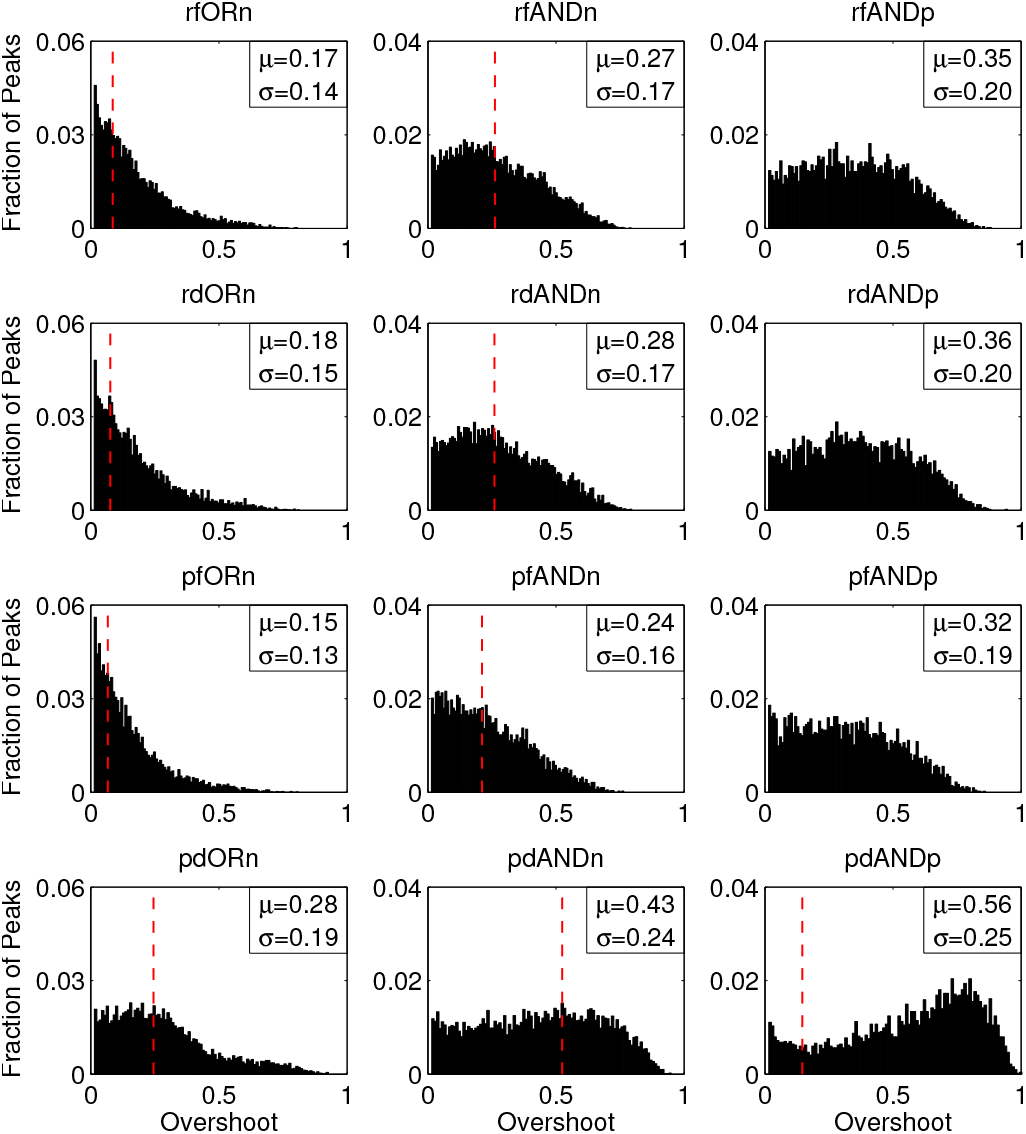
Overshoot distributions for the different composite motifs. *µ* and *σ* indicate the mean and standard deviation of the corresponding distributions. The dashed vertical red line denotes the value of overshoot for the basal parameter set, when it shows peak.

**Figure 10:**
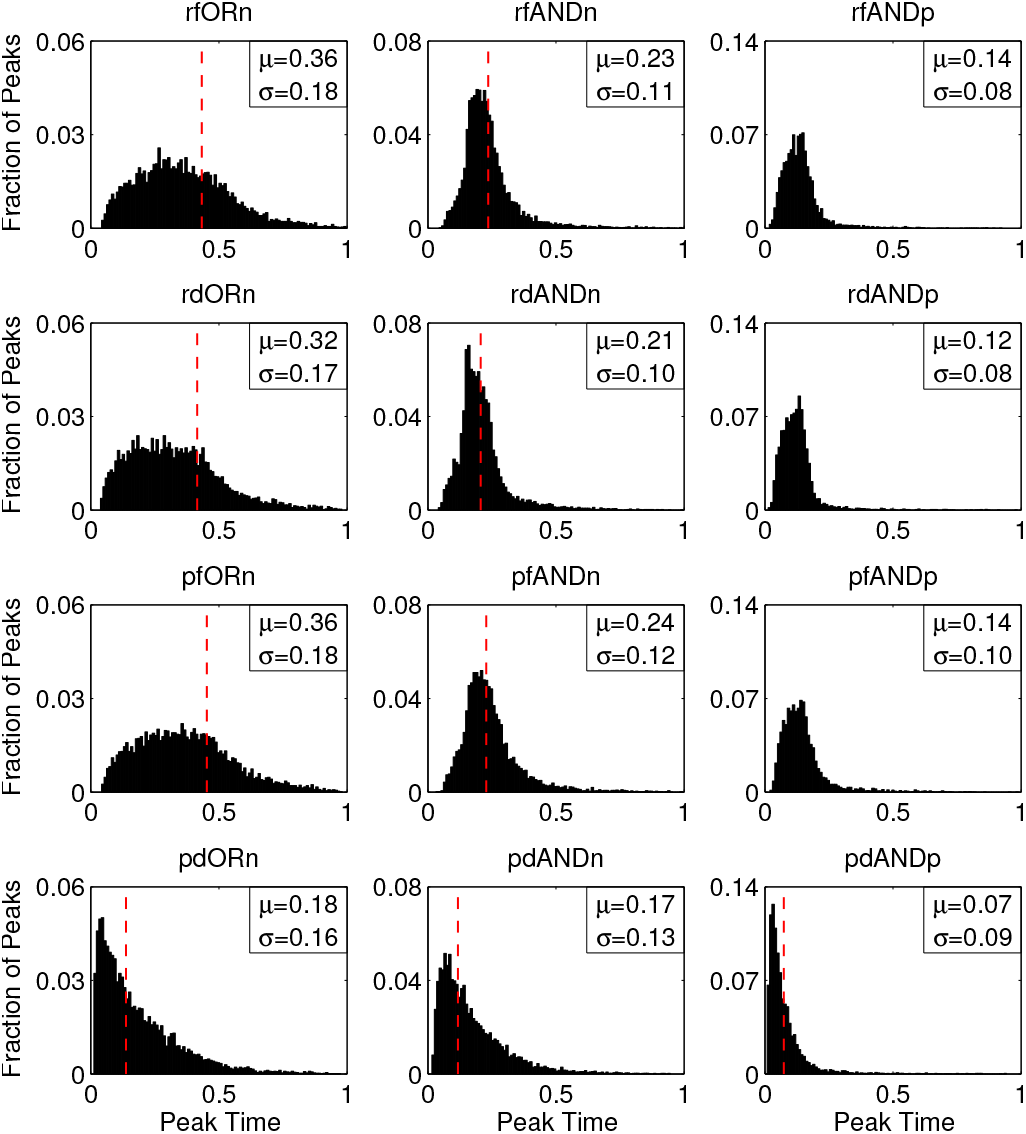
Peak time distributions for the different composite motifs. *µ* and *σ* indicate the mean and standard deviation of the corresponding distributions. The dashed vertical red line denotes the value of peak time for the basal parameter set, when it shows peak.

Regulation by protein degradation showed higher mean overshoot than the other modes of regulation. This was, however, also the case with uncoupled feedforward and feedback motifs (Fig S4). Conversely, protein degradation mediated uncoupled feedback motifs showed lower mean peak time than to other modes of regulation. However, this difference was not significant for the feedforward motifs (Fig. S5).

The distribution of peak duration did not differ significantly between the different composite motifs.

### Overshoot decreases, while peak duration changes concavely, with peak time

To understand the relationship between different peak properties I analysed how overshoot and peak duration correlated with peak time using scatter plots, for the different parameter sets that produce peaks. As in case of uncoupled feedback and feedforward motifs, overshoot correlated negatively with peak time. However, peak duration first increased and then decreased with increasing peak time suggesting that peak duration is a concave (superlinear) function of peak time. Again, this is in agreement with what we observed in case of uncoupled feedback and feedforward motifs. Considering all this, it appears that both these relationships between peak properties are some kind of a universal feature of peak generating motifs in biological systems.

To understand the reason behind this relationship, analysed a simple model of feedforward loop which has just two variables – input and controller (Sup. Sec. 4). The analysis reveals that differences in the degradation rates of output and controller is necessary for the above mentioned relationships to hold. In the absence of such a condition the peak time depends on only one parameter – the fold difference between the inputs, *I*_1_ and *I*_2_. To check if difference in degradation rates is sufficient for the observed pattern of correlation between overshoot, peak duration and peak time, I repeated the simulations with parameter sets that just varied in the degradation rate constants (1/5 to 5 times the basal value), for both uncoupled feedback and feedforward motifs, and the composite motifs. In case of the uncoupled feedback and feedforward motifs, variation in the degradation rate constants of all the species (proteins and RNA) was found to be sufficient to generate the pattern, however, variation of just the degradation rate constants of the RNA species was not sufficient (Fig. S6–S7). For the composite motifs, variation of the degradation rate constants of just the RNA speceis as well as that of all the species, generated correlation patterns between the three peak properties, showed similar trends (Fig. S8–S11).

**Figure 11:**
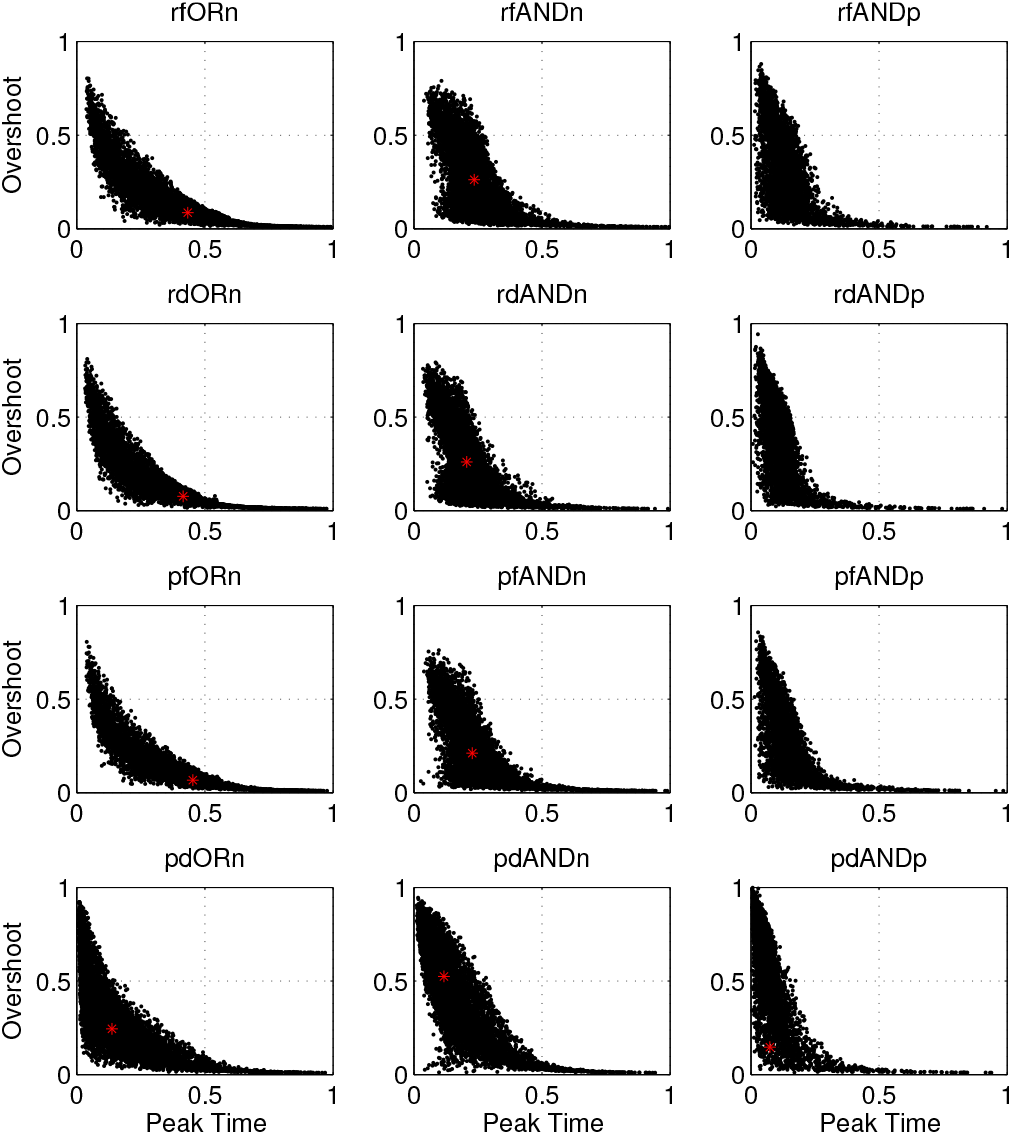
Scatter plots between peak time and overshoot for the composite motifs. Asterisks denote the values corresponding to the basal parameter set, when it shows peak.

**Figure 12:**
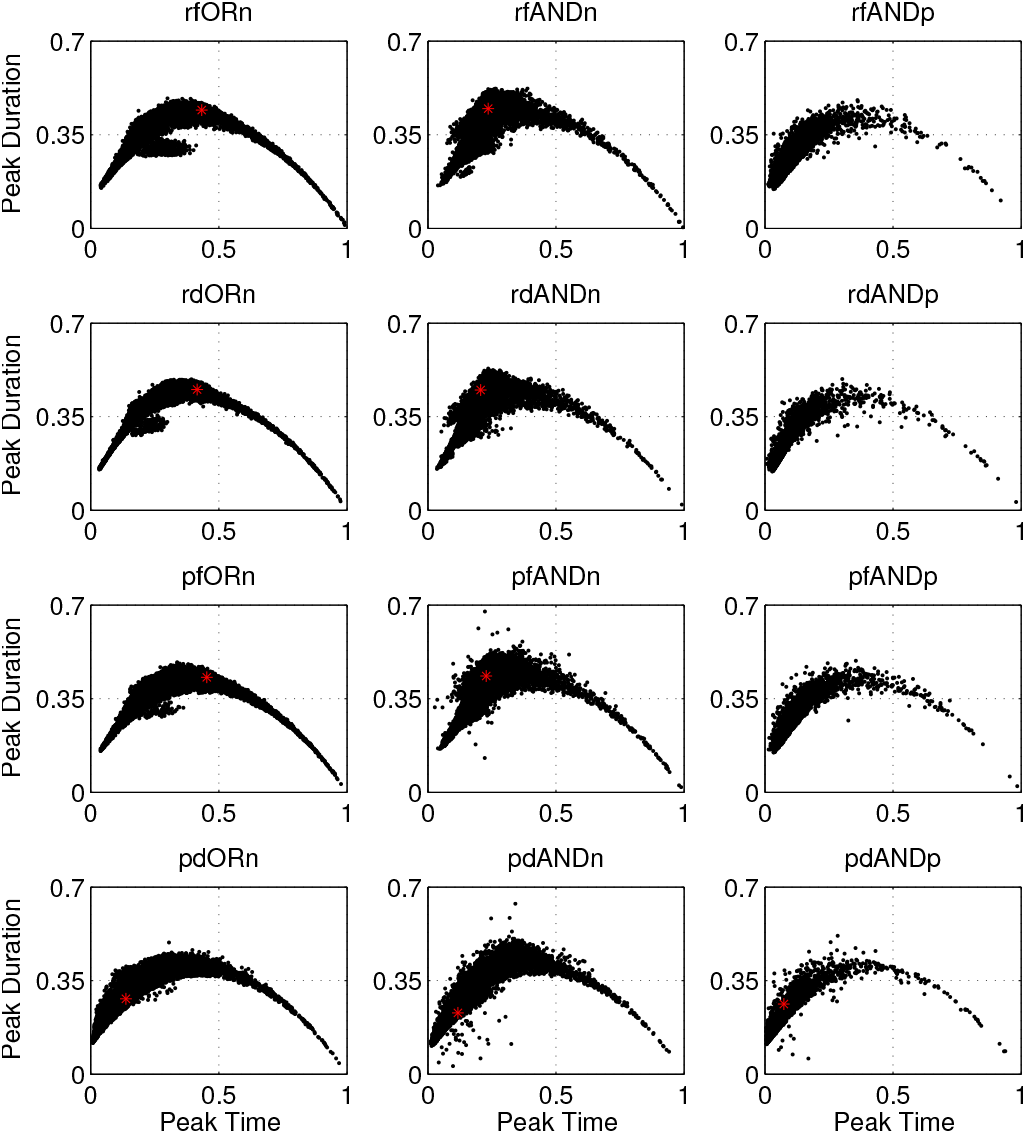
Scatter plots between peak time and peak duration for the composite motifs. Asterisks denote the values corresponding to the basal parameter set, when it shows peak.

## Conclusion

Though, feedback and feedforward motifs have been studied independently, a system that involves a coupled feedback and a feedforward motif has not been studied. I have mentioned a few examples, in the introduction ^[12, 15, 16]^ but a detailed survey of such motifs is essential. These motifs also represent a situation in which a feedback system is tuned by a common upstream regulator.

In this study, I have used a simple model to analyse the properties of different classes of composite feedforward-feedback motifs, operating via different modes of regulation. I have used a framework to simultaneously study multiple systems using a shared parameter set. This framework allows for parameter variation in order to study global sensitivity of different system properties to different parameters.

ANDn motifs show perfect adaptation (zero gain) for all modes of regulation, for most parameter sets whereas for ANDp motifs, the gain was hypersensitive to parameters such that it either showed perfect adaptation or no regulation at all. Moreover, in the ANDp motifs, the controller and output showed a reciprocal response both with respect to the steady state and the dynamics. Such motifs can switch between two contrasting regimes with slight parameter changes.

Early saturation of the input-output characteristics of ANDn motifs suggest that their adaptability is robust to a broad range of input. However, the hypersensitivity of output at low levels of input may be a trade-off. ANDp motifs also showed saturation and non-monotonicity and in some cases the characteristics were not very hypersensitive to input. This would allow these motifs to exhibit different kinds of responses to different levels of input. Such a feature may be useful in conditions where excessive stimulation by a signal would be undesirable (for example, stimulation of neurons).

The peak properties i.e. overshoot, peak time and peak duration seem share a relationship which seems to be universal for incoherent feedforward motifs, negative feedback motifs and composite motifs made of coupled feedback and feedforward loops; overshoot decreases with peak time whereas peak duration varies concavely with it. Difference in the degradation rates of the participating species, seems to be necessary and sufficient for this relationship to hold. It is important to check if other peak-generating motifs also exhibit this property.

Overall, it is evident that regulation by protein degradation is very distinct from other modes of regulation irrespective of the motif structure (even for uncoupled feedforward and feedback motifs); it has faster response and higher peaks. However, regulation by protein degradation is likely to be more expensive for the cell than the other modes of regulation because of the additional cost of translation.

## Acknowledgments

I thank CSIR-National Chemical Laboratory for providing me with resources required for conducting this study.

